# Mining the UniProtKB/Swiss-Prot database for antimicrobial peptides

**DOI:** 10.1101/2024.05.24.595811

**Authors:** Chenkai Li, Darcy Sutherland, Ali Salehi, Amelia Richter, Diana Lin, Sambina Islam Aninta, Hossein Ebrahimikondori, Anat Yanai, Lauren Coombe, René L. Warren, Monica Kotkoff, Linda M.N. Hoang, Caren C. Helbing, Inanc Birol

## Abstract

The ever-growing global health threat of antibiotic resistance is compelling researchers to explore alternatives to conventional antibiotics. Antimicrobial peptides (AMPs) are emerging as a promising solution to fill this need. Naturally occurring AMPs are produced by all forms of life as part of the innate immune system. High-throughput bioinformatics tools have enabled fast and large-scale discovery of AMPs from genomic, transcriptomic, and proteomic resources of selected organisms. Public protein sequence databases, comprising over 200 million records and growing, serve as comprehensive compendia of sequences from a broad range of source organisms. Yet, large-scale *in silico* probing of those databases for novel AMPs has never been reported. In the present study, we propose an AMP mining workflow to predict novel AMPs from the UniProtKB/Swiss-Prot database using the AMP prediction tool, AMPlify, as its discovery engine. Using this workflow, we identified 8,008 novel putative AMPs from all eukaryotic sequences in the database. Focusing on the practical use of AMPs as suitable antimicrobial agents with applications in the poultry industry, we prioritized 40 of those AMPs based on their similarities to known chicken AMPs in predicted structures. In our tests, 13 out of the 38 successfully synthesized peptides showed antimicrobial activity against *Escherichia coli* and/or *Staphylococcus aureus*.

## Introduction

As a consequence of the worldwide overuse of antibiotics, the world is under the threat of entering a “post-antibiotic era” [1]. The decreasing effectiveness of conventional antibiotics is posing great challenges in the treatment of many infectious diseases [1], with an estimated 1.27 million people dying due to antibiotic resistance in 2019 [2].

Aside from the extensive use of conventional antibiotics in clinical settings, antibiotics are also widely used in the agriculture industry [3]. Certain multidrug-resistant (MDR) bacteria can be transmitted between humans and other animals, which further exacerbates the problem [3]. While the occurrence of antibiotic resistance is increasing, there has been a substantial decline in the discovery of new antimicrobial agents since the 1990s [4,5]. Consequently, there is an urgent need for the discovery of novel and effective substitutes for conventional antibiotics.

Antimicrobial peptides (AMPs), a family of short and often cationic peptides, are regarded as one promising substitute for conventional antibiotics [6]. Naturally occurring AMPs are produced by all life forms as part of the innate immunity [7], usually in an inactive precursor form with a signal peptide, an acidic pro-sequence, and the bioactive mature peptide [8]. The bioactive mature AMPs are released by proteolytic cleavage of their precursors [7]. While the majority of known AMPs recorded in public AMP databases have been shown to possess antibacterial activity [9], many of them have also been reported with other types of antimicrobial activities, including antifungal [10] and antiviral [11]. Most AMPs exert their effects by directly interacting with bacterial membranes or cell walls, causing non-enzymatic disruption [7]. Additionally, some eukaryotic AMPs also perform modulation of immune responses [7,12]. In comparison with conventional small-molecule antibiotics, which have specific functional or structural targets, the diverse modes of action of AMPs may hold an advantage in being able to better overcome bacterial resistance [13]. Nevertheless, it is still possible to observe resistance to AMPs if bacteria are exposed to AMPs for extended periods of time [13], highlighting a pressing need to augment our peptide-based therapeutics arsenal.

Traditional approaches of discovering naturally occurring AMPs through wet lab screening are time-consuming, labor-intensive, and costly [14]. In the past few decades, a series of machine learning based high-throughput *in silico* AMP prediction tools have been developed to overcome this problem, including iAMP-2L [15], iAMPpred [16], AMP Scanner Vr.2 [17], and AMPlify [18,19]. Among those tools, AMPlify, which uses a deep learning model with attention mechanisms [20,21], has become the state-of-the-art method for AMP prediction. It has also successfully identified over a hundred lab-validated AMPs from amphibian and insect genomes and transcriptomes [18,22,23].

Current AMP mining studies mostly focus on genomic [18,24–26], transcriptomic [22,23], or proteomic [27] resources, with specific organisms chosen for analysis. AMP mining by large-scale scanning of public protein sequence databases, such as UniProt [28] (https://www.uniprot.org), in which the majority of the sequences have already been annotated with their functions, has never been reported. We note that mining AMPs using comprehensive protein sequence databases offers another valuable strategy for discovering novel AMPs. First, these large protein sequence databases allow researchers to discover a broad source of novel AMPs. Second, there are many uncharacterized sequences awaiting annotation in public databases, which represent an untapped source of AMPs. Further, it would be of interest to uncover possible antimicrobial properties of characterized proteins or peptides, where only non-antimicrobial functions have been reported to date. Studies have found that some proteins or peptides with other biological functions exhibit antimicrobial properties [29,30]. Histatins, for example, a series of peptides found in saliva, belong to this category. Histatins harbor multiple functions beneficial to oral health besides antimicrobial activity, including oral hemostasis, development of acquired tooth pellicle, and assistance in bonding of some metal ions [30].

Herein, we present an AMP mining workflow with AMPlify as the core AMP prediction module to predict novel AMPs from all eukaryotic sequences in the UniProtKB/Swiss-Prot database [28], which contains only manually annotated and reviewed records from the larger UniProt database. With this workflow, we identified 8,008 novel putative AMPs. Motivated by the growing global concern about the transmission of antibiotic-resistant bacteria from other animals to humans – a direct consequence of antibiotics overuse in animal agriculture, including poultry farming [31] – we conducted *in vitro* testing on a total of 38 predicted peptides. The selection of this set of peptides was based on their similarities to known chicken AMPs in predicted 3D structures.

We conducted our tests with the synthesized peptides using lab strains of *Escherichia coli* ATCC 25922 and *Staphylococcus aureus* ATCC 29213. We observed that 13 of them demonstrated antimicrobial activity against at least one bacterial strain. Chicken AMPs have theoretically been evolved to fight against pathogens in their environment [7], hence exogenous AMPs with high structural similarities to known chicken AMPs may help them cope with bacterial challenges in the farm environment.

## Results

### Integration of AMPlify balanced and imbalanced models

The AMP prediction module of the AMP mining workflow utilizes two AMPlify models, balanced and imbalanced [18,19], to obtain a curated set of putative AMP sequences. The balanced model of AMPlify exhibited superior performance on highly curated candidate sequence sets characterized by lower noise and higher confidence levels, as evidenced by its success in mining diverse genomes from the bullfrog and others [18,19,22,23]. In contrast, running AMPlify using an imbalanced model has been shown to effectively handle large, highly imbalanced candidate sequence sets enriched with non-AMPs [19]. Based on the performance of the two models in their respective advantageous application scenarios, we incorporated both in the reported AMP mining workflow. As shown in **Supplementary Figure S1**, the prediction module (Steps 3 and 4) performed a two-stage filtering by applying the imbalanced and balanced models in turn. The integration of the two models can also be considered as a filtering scheme of only selecting sequences that are predicted by both models as AMPs.

In our previous work, we built two test sets to evaluate the performance of the balanced and imbalanced models [19]. The balanced test set comprises 835 AMPs and 835 non-AMPs, while the imbalanced test set comprises 835 AMPs and 25,689 non-AMPs [19]. **Table 1** shows the performance comparison of the balanced and imbalanced models of AMPlify as well as the integration of the two (noted as “balanced + imbalanced”) on the two test sets with regard to accuracy, sensitivity, specificity, F1 score and area under the receiver operating characteristic curve (AUROC). The integrated model outputs predicted labels instead of probabilities, hence the AUROC values were not reported for it.

**Table 1:** Performance comparison of the balanced/imbalanced models of AMPlify, as well as the integration of the two, on the balanced and imbalanced test sets. Values of accuracy (acc), sensitivity (sens), specificity (spec), F1 score (F1) and area under the receiver operating characteristic curve (AUROC) are presented in percentages.

| **AMPlify test set** | **AMPlify model** | **Acc** | **Sens** | **Spec** | **F1** | **AUROC** |
| --- | --- | --- | --- | --- | --- | --- |
| balanced | balanced | **93.71** | 92.93 | 94.49 | **93.66** | **98.37** |
|  | imbalanced | 89.22 | **94.37** | 84.07 | 89.75 | 96.88 |
|  | balanced + imbalanced<sup>a</sup> | 93.59 | 91.50 | **95.69** | 93.46 | —* |
| imbalanced | balanced | 95.97 | 92.93 | 96.06 | 59.19 | 98.06 |
|  | imbalanced | 98.94 | **94.37** | 99.09 | 84.87 | **99.54** |
|  | balanced + imbalanced<sup>a</sup> | **99.31** | 91.50 | **99.56** | **89.25** | —* |

<sup>a</sup> Integration of the two models: only sequences predicted as AMPs by both models are determined to be AMPs.
\* AUROC values were not calculated for models which only output predicted labels instead of probabilities.

On the balanced test set, where the AMPlify balanced model performs better than the imbalanced model except for sensitivity [19], the integrated model does not show much improvement when compared with using the two models independently, with only an increase in specificity by 1.2% (95.69% vs. 94.49%).

On the imbalanced test set, where the AMPlify imbalanced model performs better than the balanced model in all five metrics [19], the integrated model achieves improvement in accuracy (99.31% vs. 98.94%), specificity (99.56% vs. 99.09%), and F1 score (89.25% vs. 84.87%). For highly imbalanced candidate sequence sets where non-AMP sequences far outnumber AMP sequences, it is crucial to minimize the number of false positives to prevent an excessive number of non-AMP sequences from being forwarded to downstream *in vitro* validation. In this case, although there is a slight decrease in sensitivity (91.50% vs. 94.37%), the improvement in other metrics, particularly specificity, suggests that the integrated model performs better than the imbalanced model alone under this scenario.

Additionally, all AMPlify models (balanced, imbalanced, and their integration) outperform other existing AMP prediction methods for comparison on the two test sets (**Supplementary Table S1**, **Supplementary Table S2**).

Considering that our source for AMP mining is a large protein sequence database [28], in which most of the sequences are not expected to be AMPs, filtering for putative AMP sequences by integrating balanced and imbalanced models is a sensible way to reduce the number of false positives.

### Predicted AMPs

By applying the AMP mining workflow to all eukaryotic sequences in UniProtKB/Swiss-Prot [28], 10,720 distinct candidate mature peptide sequences were predicted as AMPs, of which 8,008 (74.70%) were novel putative AMPs (**Supplementary Figure S1**). All parent sequences were cleaved by the cleavage module of AMP discovery pipeline rAMPage v1.0.1 [22], which adapts ProP v1.0c [32], for mature peptide sequences. Sequences without predicted signal peptides were filtered out in Step 2 of the workflow (**Supplementary Figure S1**).

The 8,008 novel putative AMPs have an average length of 52.49 aa (SD = 26.49 aa; aa: amino acids; SD: standard deviation) and an average net charge of 3.04 (SD = 3.88), while the 4,538 known AMP sequences have an average length of 30.21 aa (SD = 20.28 aa) and an average net charge of 3.05 (SD = 3.10) (**Supplementary Figure S2**). A one-sided Welch’s t-test indicates that the larger average length of the novel putative AMPs compared with that of the known AMP sequences is statistically significant (p < 0.05). However, researchers tend to prioritize shorter sequences for validation due to their lower synthesis costs for *in vitro* validation [22], which could explain the difference in average lengths. The novel putative AMP sequences also exhibit a notable level of novelty, sharing a low sequence similarity level of 32.71%, on average, to the known AMP sequences (**Figure 1**).

**Figure 1:**
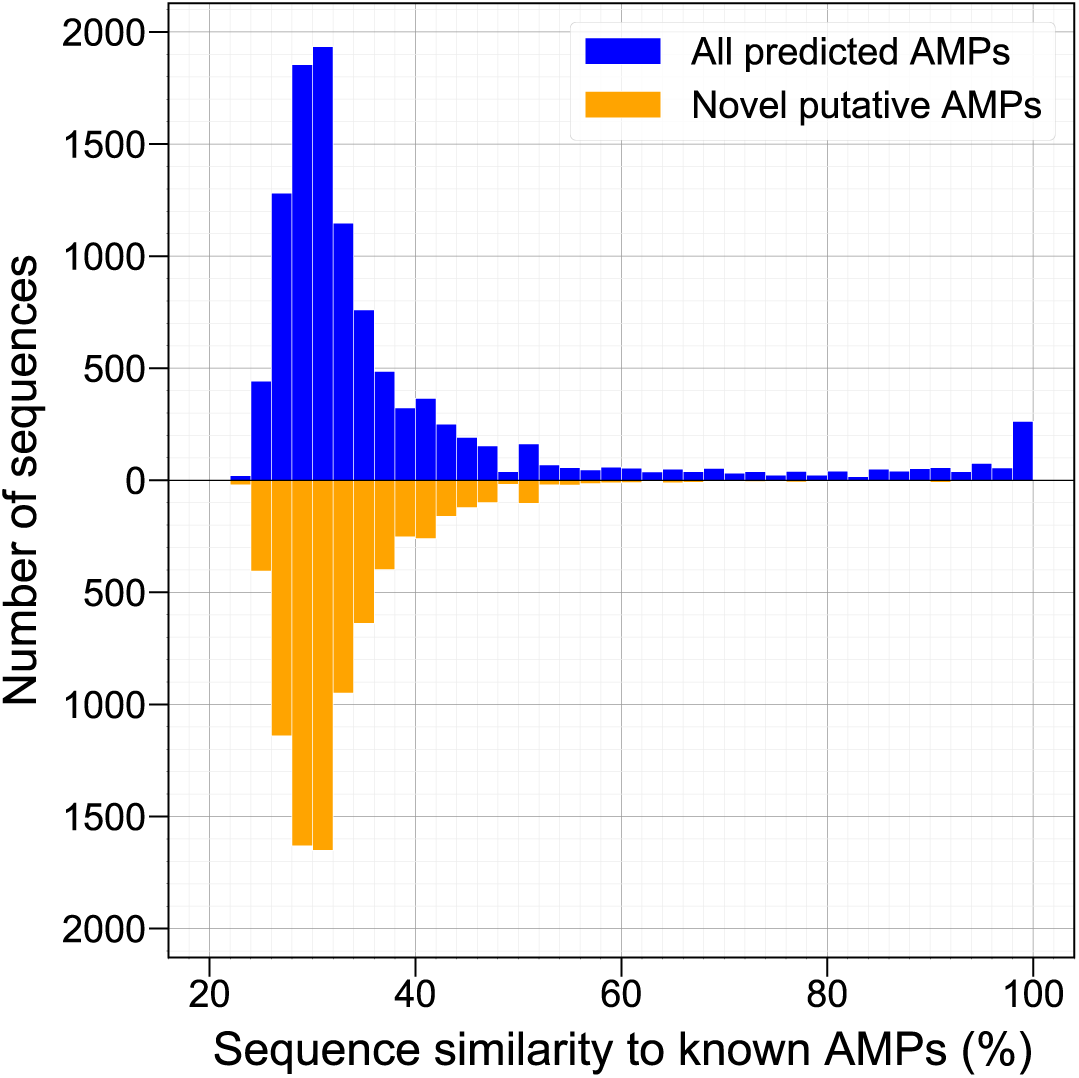
Sequence similarity distributions of the predicted AMPs from the UniProtKB/Swiss-Prot database to known AMPs. The sequence similarity distribution of all 10,720 predicted AMPs to known AMPs from Antimicrobial Peptide Database (APD3) [9] and Database of Anuran Defense Peptides (DADP) [33] was visualized, along with that of the 8,008 novel putative AMPs from all predicted AMPs. The former distribution holds a mean of 38.27% and a standard deviation of 17.28%, while the latter holds a mean of 32.71% and a standard deviation of 7.21%. The sequence similarity of each predicted AMP to known AMPs was considered as the sequence similarity of that predicted AMP sequence to its most similar sequence in the known AMP set, based on which the distributions were plotted.

Tracing back to the original source entries in the protein sequence database, the 10,720 predicted AMPs corresponded to 8,481 parent sequences from a total of 8,862 UniProt entries. In contrast, the 8,008 novel putative AMPs corresponded to 6,349 parent sequences from 6,654 UniProt entries (**Supplementary Figure S1**). In our analysis, we define parent sequences with AMPs predicted from their cleaved mature peptide sequences to be putative AMP precursor sequences, and UniProt entries of those putative AMP precursor sequences to be putative AMP entries.

Among all 8,481 putative AMP precursor sequences identified, 86.84% (7,365) only had one distinct predicted AMP sequence, as shown in **Figure 2**. However, we did notice cases where multiple AMP sequences were predicted from a single putative precursor sequence [33]. Specifically, 698 putative precursor sequences were predicted with two distinct AMPs from each, 207 predicted with three, 74 predicted with four, and 57 predicted with five. There were 80 putative precursor sequences with more than five distinct AMPs predicted from each.

**Figure 2:**
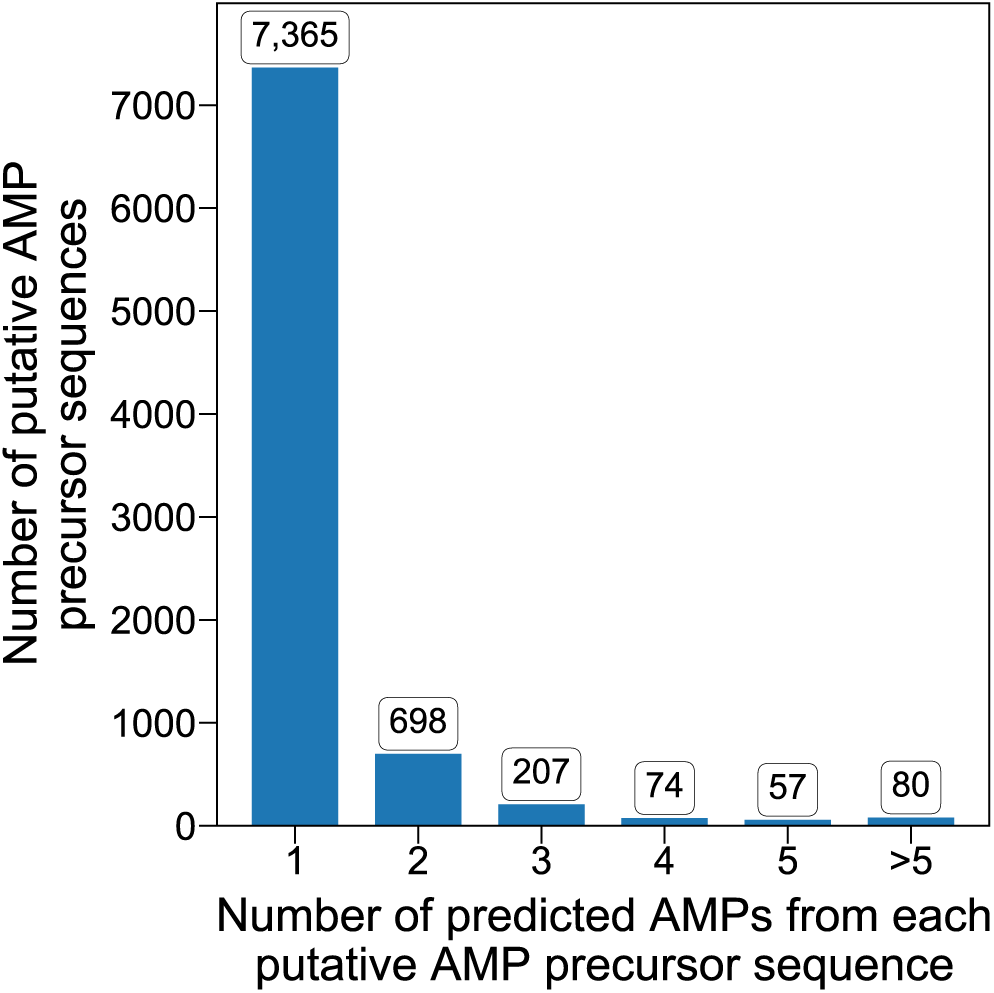
Distribution for the number of predicted mature AMP sequences found within each putative AMP precursor sequence mined from the UniProtKB/Swiss-Prot database. The bar chart was plotted based on 8,481 distinct putative AMP precursor sequences from the 8,862 putative AMP entries. Each bar shows the number of putative AMP precursor sequences that were predicted with the corresponding number of distinct mature AMPs by AMPlify.

The doughnut chart in **Figure 3** categorizes the 8,862 putative AMP entries identified by the AMP mining workflow by source organisms. According to the statistics reported by the Antimicrobial Peptide Database (APD3, https://aps.unmc.edu) web server [9] on January 2023, out of all 3,569 sequences in records, amphibians are the largest organism source of AMPs (1,196 sequences), followed by bacteria (380), plants (371), insects (367), and mammals (363) [9]. Based on this, all the putative AMP entries we report herein were classified into the following five categories: amphibian, plant, insect, mammalian, and other entries. The category of bacterial AMPs was not included, as we only considered eukaryotic AMPs in this work.

**Figure 3:**
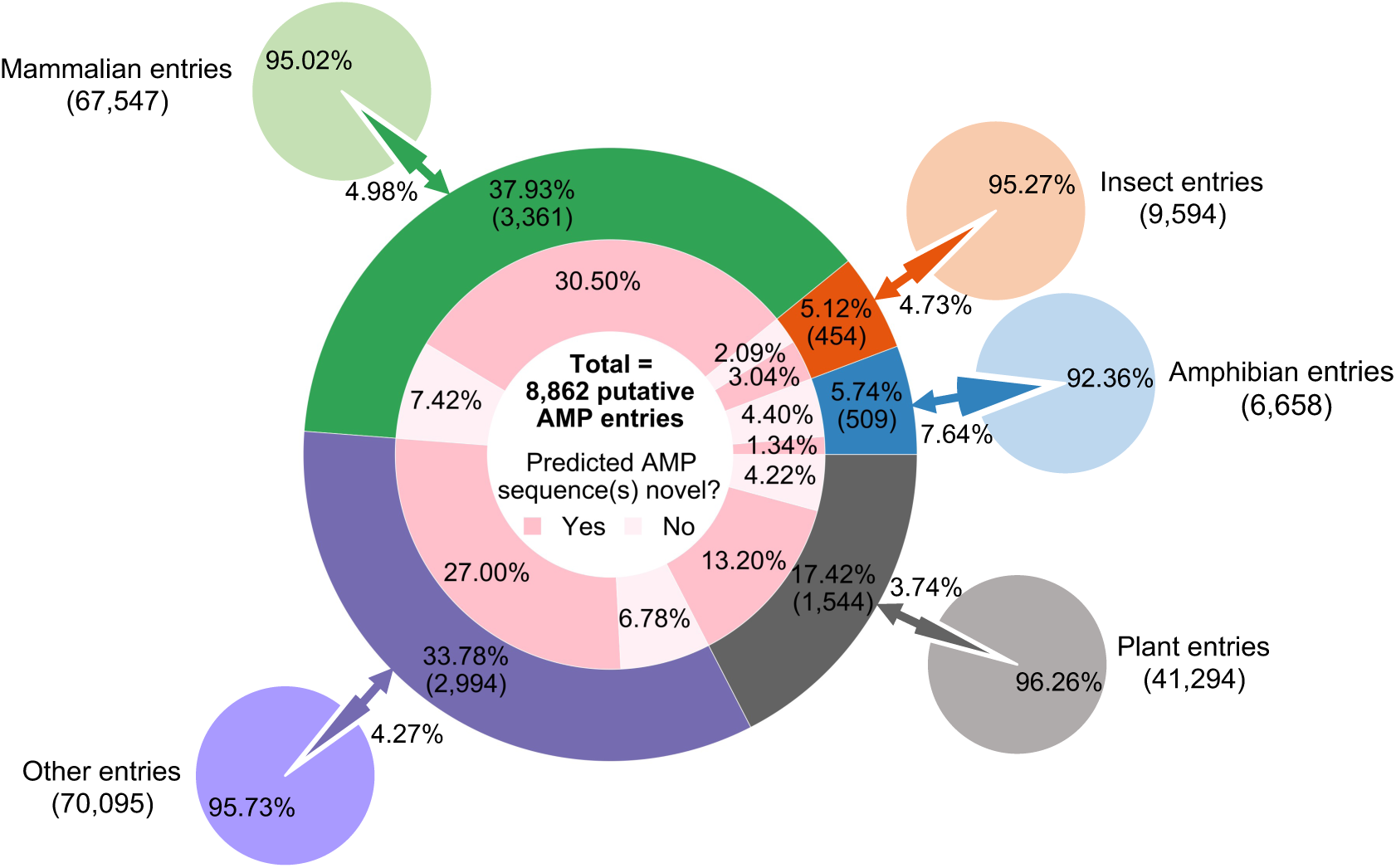
Categorization of the putative AMP entries identified from the UniProtKB/Swiss-Prot database based on source organisms. The 8,862 putative AMP entries with mature AMPs predicted by AMPlify were classified into five categories (mammalian, plant, amphibian, insect, and other entries) based on their source organisms, as shown in the middle doughnut chart. The predicted AMPs were further checked against the known AMP sequences in Antimicrobial Peptide Database (APD3) [9] and Database of Anuran Defense Peptides (DADP) [33] as well as the annotations of the corresponding UniProt entries. Those not found in those two AMP databases and not annotated with the AMP-related keywords were labeled as novel AMP sequences. The five small pie charts surrounding the middle doughnut chart provide additional information on the percentages of entries in UniProtKB/Swiss-Prot identified as putative AMP entries by the AMP mining workflow.

The distribution of source organisms for putative AMP entries is highly variable, reflecting the composition of the UniProtKB/Swiss-Prot database (**Figure 3**). Mammalian entries constitute the largest source organism category for putative AMP entries, comprising 37.93% (3,361/8,862) of all. Out of all 3,361 putative AMP entries from the mammalian category, 2,703 (80.42%) of them had novel AMPs predicted from their cleaved candidate mature peptide sequences, making it the largest source organism category for novel putative AMP entries as well. Putative AMP entries not belonging to any of the four specified source organism categories also make up a large portion, following the mammalian category, and represent 33.78% (2,994/8,862) of all putative AMP entries. In this category of putative AMP entries, 79.93% (2,393/2,994) had novel AMPs predicted from their cleaved candidate mature peptide sequences. Plant entries form the third largest group among all putative AMP entries, contributing 17.42% (1,544/8,862) to the total. Overall, 1,170 out of 1,544 putative plant AMP entries (75.78%) were considered as novel discoveries. Amphibian and insect categories are the two smallest in this analysis, with 509 and 454 putative AMP entries in each. While 59.25% (269/454) of the putative insect AMP entries were determined to be novel discoveries, the amphibian category is the only source organism category with more than half of the putative AMP entries (76.62%) already identified with known mature AMP sequences and/or annotated with AMP-related keywords. We postulate the main reason for this is that amphibians are considered as a rich source of AMPs [34], hence has become a popular source for AMP mining [18,22,23], resulting in more of the amphibian AMPs having been discovered and annotated than those from other organisms. Moreover, amphibians are also the source organism group with the largest proportion of entries (7.64%) in UniProtKB/Swiss-Prot [28] identified as putative AMP entries by the AMP mining workflow, followed by mammals (4.98%), insects (4.73%), source organisms other than the specified ones (4.27%), and plants (3.74%), as can be seen from the pie charts in **Figure 3**.

We did not observe any instances where a single putative AMP precursor sequence had novel putative AMP(s) predicted, while also containing known AMP(s) in its cleaved candidate mature peptides and/or being annotated with AMP-related keywords (described in the **Materials and methods** section). We also note that UniProt entries annotated with AMP-related keywords can be proteins or peptides that assist in the defense against microbes rather than possessing antimicrobial activities themselves. However, to ensure a cleaner set of novel putative AMPs, all the predicted AMPs with their parent sequence entries annotated with AMP-related keywords were not considered as novel discoveries.

### *In vitro* validation results

Considering the large number of putative AMPs identified by the AMP mining workflow, we prioritized a subset of the sequences for synthesis and *in vitro* validation. In this work, we primarily focused on the applications of AMPs in poultry farming. Since three-dimensional (3D) structures of proteins or peptides may be a determining factor for their biological functions [35], we prioritized putative AMPs based on their similarities to known chicken AMPs in 3D structures predicted by ColabFold [36]. The reference chicken AMP for a putative AMP was defined as the most similar known chicken AMP to that putative AMP in predicted 3D structures.

We selected a total number of 40 short cationic novel putative AMP sequences with AMPlify scores ≥ 10 for synthesis based on their similarities to known chicken AMPs in predicted structures, as measured by template modeling scores (TM-scores) [37,38] (**Supplementary Figure S3**). AMPlify score is an AMP prediction score reported by AMPlify [18,19], and TM-score is a protein structural similarity score reported by TM-align [38] (see the **Materials and methods** section for details). Of those selected, 30 had the highest TM-scores regarding their similarities to known chicken AMPs in predicted structures, while the remaining 10 had the lowest TM-scores. **Supplementary Table S3** and **Supplementary Table S4** list the characteristics of the 40 peptides and their original UniProt entry information, respectively. We note that two of the top 30 sequences were not successfully synthesized, resulting in a final list of 38 putative AMPs for *in vitro* validation. The three reference chicken AMPs, Chicken CATH-2 [39], Chicken CATH-3 [40], and Ovipin [41], that matched the top 30 sequences were also tested for comparison (**Supplementary Table S5**).

All synthesized peptides were tested against two bacterial isolates: the Gram-negative *E. coli* ATCC 25922, and the Gram-positive *S. aureus* ATCC 29213. Porcine red blood cells (RBCs) were used to assess the hemolytic activity of the peptides. In this study we used porcine RBCs, as observations in prior work confirmed that porcine and chicken RBCs report similar results in hemolytic activity measurements (data not shown). Out of the 38 putative AMPs synthesized, 13 displayed antimicrobial activity against *E. coli* ATCC 25922, with three of the 13 additionally active against *S. aureus* ATCC 29213. **Figure 4** summarizes the antimicrobial and hemolytic activities of the 13 active peptides in minimum inhibitory concentration (MIC) and the concentration that lyses 50% of the RBCs (HC_50_), respectively, with the entire *in vitro* validation results of all peptides shown in **Supplementary Table S6**. The left section of **Figure 4** shows the results for the 11 active peptides from the set of 30 putative AMPs with highest TM-scores, as well as their reference chicken AMPs for comparison. Results of the two active peptides from the set of 10 putative AMPs with lowest TM-scores are shown in the right section of **Figure 4**. **Supplementary Figure S4** additionally shows the activity of all tested putative AMPs regarding their AMPlify scores and similarities to known chicken AMPs in predicted structures.

**Figure 4:**
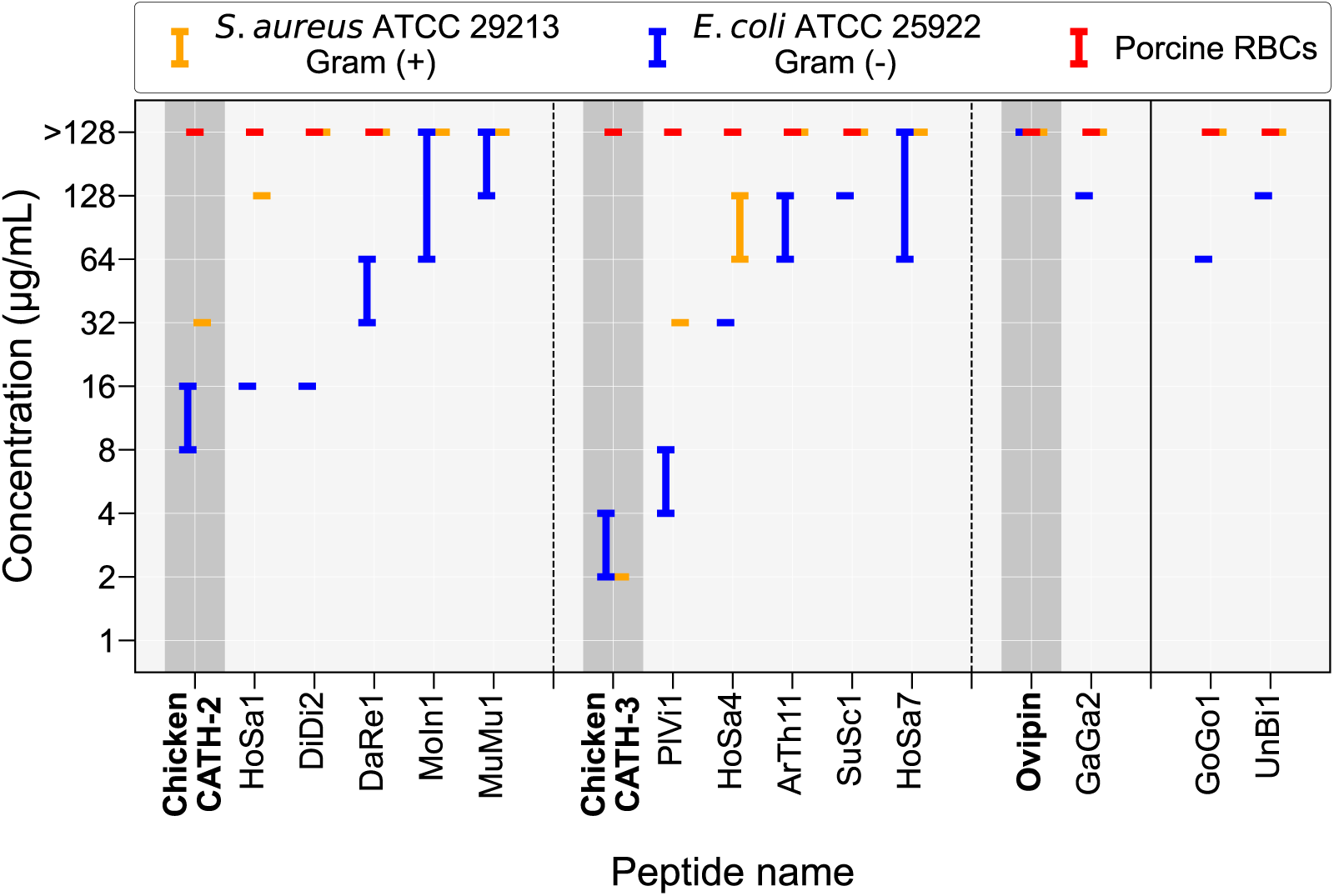
Antimicrobial and hemolytic activities of the 13 novel AMPs mined from the UniProtKB/Swiss-Prot database that were active against at least one bacterial strain of *Escherichia coli* ATCC 25922 and *Staphylococcus aureus* ATCC 29213. Antimicrobial and hemolytic activities were measured by minimum inhibitory concentration (MIC) and concentration that lyses 50% (HC_50_) of the red blood cells (RBCs), respectively. HC_50_ was determined using porcine RBCs. Data is presented as the lowest effective peptide concentration range (μg/mL) observed in three independent experiments performed in duplicate, with one maximum data point and one minimum data point dropped for each measurement. The results are divided into two sections, as separated by the solid vertical line. The left section shows results of the 11 active peptides that were most similar to known chicken AMPs in predicted three-dimensional (3D) structures as measured by TM-scores, while the right section shows results of the two active peptides that were least similar to known chicken AMPs in predicted 3D structures. These 11 active peptides in the left section are categorized into three sub-sections, as separated by the dashed vertical lines, according to their reference chicken AMPs (i.e., the most similar known chicken AMP to each putative AMP in predicted 3D structures). The results of the three reference chicken AMPs: Chicken CATH-2, Chicken CATH-3, and Ovipin (dark grey background), are listed for comparison in each sub-section. We note that hemolysis experiments were not performed for putative AMPs that did not show any antimicrobial activity (MIC > 128 μg/mL) in at least two repeats for each bacterial strain tested (i.e., MoIn1, MuMu1, and HoSa7).

Among the 10 tested putative AMPs with Chicken CATH-2 as their reference chicken AMP, five showed antimicrobial activity in our tests. Within this group of five peptides, HoSa1, with its original UniProt entry annotated as an olfactory receptor from humans [28], possessed the strongest antimicrobial activity against *E. coli* ATCC 25922 (MIC = 16 μg/mL), and was the only peptide that showed additional activity against *S. aureus* ATCC 29213 (MIC = 128 μg/mL). DiDi2 showed the same antibacterial activity as HoSa1 against *E. coli* ATCC 25922 with an MIC of 16 μg/mL, with its original UniProt entry annotated as a putative uncharacterized transmembrane protein from social amoebas [28]. DaRe1, the parent sequence of which is annotated as a GTP-binding protein from zebrafish, inhibited the growth of *E. coli* ATCC 25922 at an MIC of 32–64 μg/mL. MoIn1 and MuMu1 presented the weakest antibacterial activity against *E. coli* ATCC 25922 in this group, with MICs of 64–>128 μg/mL and ≥128 μg/mL. The parent sequence of MoIn1 is annotated as part of an intermediate translocation complex at the inner envelope membrane of chloroplasts from mulberries, while the parent sequence of MuMu1 is annotated as a mouse surfeit locus protein, a component of the MITRAC (mitochondrial translation regulation assembly intermediate of cytochrome c oxidase complex) [28]. The reference chicken AMP, Chicken CATH-2, possessed stronger antimicrobial activity against *E. coli* ATCC 25922 (MIC = 8–16 μg/mL) and *S. aureus* ATCC 29213 (MIC = 32 μg/mL) than all five peptides in this group.

Among the 16 tested putative AMPs with Chicken CATH-3 as their reference chicken AMP, five showed antimicrobial activity in our tests. Among the five peptides in the group, PlVi1 had the strongest antimicrobial activity against both *E. coli* ATCC 25922 (MIC = 4–8 μg/mL) and *S. aureus* ATCC 29213 (MIC = 32 μg/mL), and it was also the most active peptide among all 38 putative AMPs tested. The parent sequence of PlVi1 is annotated as the precursor of a RxLR effector protein that completely suppresses the host cell death induced by cell death-inducing proteins, which is derived from a type of oomycete that causes the downy mildew disease of grapevines [28]. HoSa4, derived from a human IQ domain-containing protein without a detailed functional annotation [28], was the second most active AMP in this group. HoSa4 was observed to be bioactive against *E. coli* ATCC 25922 and *S. aureus* ATCC 29213 with MICs of 32 μg/mL and 64–128 μg/mL, respectively. We note that PlVi1 and HoSa4 are the only two peptides that were active against both bacterial strains tested in this group. ArTh11 inhibited the growth of *E. coli* ATCC 25922 at an MIC of 64–128 μg/mL. The parent sequence of ArTh11 is annotated as an F-box protein from mouse-ear cress [28], with no specific functional annotation. SuSc1 and HoSa7 were the two least active peptides in the group, with bioactivity against *E. coli* ATCC 25922 at MICs of 128 μg/mL and 64–>128 μg/mL, respectively. SuSc1 was derived from a precursor sequence of adrenomedullin in pigs and cattle, while HoSa7 was cleaved from a human G-protein coupled receptor [28]. The reference chicken AMP, Chicken CATH-3, harbored stronger antimicrobial activity against *E. coli* ATCC 25922 (MIC = 2–4 μg/mL) and *S. aureus* ATCC 29213 (MIC = 2 μg/mL) than all five peptides in this group. Two peptides in the top 30 putative AMP list matched Ovipin as their reference chicken AMP, with one (GaGa2) showing antimicrobial activity in our tests. However, it only showed minimal antibacterial activity against *E. coli* ATCC 25922 (MIC = 128 μg/mL), with no activity against *S. aureus* ATCC 29213 (MIC > 128 μg/mL). This peptide was derived from a chicken tenascin [28]. The reference chicken AMP, Ovipin, did not show any activity against the two bacterial strains tested in the present study (MIC > 128 μg/mL) but was previously reported to be bioactive against a Gram-positive *Micrococcus luteus* strain [41].

Among the 10 least similar putative AMPs to known chicken AMPs in predicted structures, only two (GoGo1 and UnBi1) showed activity in our tests. GoGo1 was part of the sequence of a transcriptional repressor found in five different primate species of western lowland gorillas, bonobos, Bornean orangutans, chimpanzees, as well as humans [28]. It inhibited the growth of *E. coli* ATCC 25922 with an MIC of 64 μg/mL. The parent peptide of UnBi1 was annotated as a neurotoxin from sea snails [28]. It only showed minimal activity against *E. coli* ATCC 25922 (MIC = 128 μg/mL). Neither of these two peptides were active against *S. aureus* ATCC 29213 (MIC > 128 μg/mL).

We note that hemolysis experiments were not performed for putative AMPs that did not show any antimicrobial activity (MIC > 128 μg/mL) in at least two repeats for each bacterial strain tested, resulting in MoIn1, MuMu1, and HoSa7 not being characterized for hemolytic activity (**Supplementary Table S6**). None of the other 10 novel bioactive AMPs were hemolytic to the porcine RBCs (HC_50_ > 128 μg/mL).

## Discussion

In this work, we applied an AMP mining workflow to discover novel eukaryotic AMPs from the UniProtKB/Swiss-Prot database [28]. The AMP mining workflow utilizes the state-of-the-art AMP prediction tool AMPlify [18,19], as well as the rAMPage cleavage module [22] with ProP [32] for precursor sequence cleavage. The scripts for the AMP mining workflow have been incorporated into AMPlify v2.0.0 and are available at https://github.com/bcgsc/AMPlify. Using this workflow, we identified 8,008 distinct novel AMP sequences from all eukaryotic sequences in the UniProtKB/Swiss-Prot database. We have made all predicted AMPs in the present study publicly available through a Zenodo repository at https://doi.org/10.5281/zenodo.8133088 [42], providing the community with putative AMP sequences to validate in the lab and to extend the current arsenal of peptide-based therapeutics.

While the presented AMP mining workflow successfully identified a considerable number of putative AMPs, it is essential to acknowledge the limitations inherent in the machine learning based tools we employed. Both AMPlify and ProP, although powerful, are not infallible, as neither of them achieves 100% accuracy in their respective tasks. As is common with most machine learning based bioinformatics tools, their performance can be limited by the quality as well as the size of available data used for training [18,19]. Despite these limitations, the reported performance of AMPlify and ProP [18,19,32] still establishes them as state-of-the-art tools, rendering them highly suitable for integration into an AMP mining workflow. We anticipate that the limitations of these tools will gradually diminish as more training data become available [18] and machine learning techniques continue to advance [43].

Herein, we focused on AMPs that have potential utility in poultry farming. As the first step towards a field application, we initially tested our selected putative AMPs against *E. coli* and *S. aureus*, and found 13 to be active against at least one of the bacterial strains tested. A total of 11 of these active peptides were relatively similar to chicken AMPs in their predicted 3D structures (TM-scores ranging from 0.6934 to 0.8352), 10 of which were from organisms other than chickens. Based on the fact that the biological function of a protein or peptide can be influenced by the 3D structure it adopts [35] and chicken AMPs have evolved to fight the pathogens that infect their hosts [7], these newly discovered AMPs with high similarities to chicken AMPs in predicted structures may target similar pathogens as chicken AMPs do. As a result, we foresee the potential usage of these characterized AMPs as possible substitutes for conventional antibiotics to fight against pathogens in chicken farming.

Although some of the putative AMPs were not active against the bacterial strains we tested, they may still be active against other pathogens, especially those that infect chickens. Further tests on a broader range of bacterial species could be done to validate their application as novel antibiotics for chickens.

While it is hypothesized that AMPs may not induce antibiotic resistance to the extent of conventional antibiotics [13], cases of bacterial cross-resistance to multiple AMPs have been reported [44], raising a concern regarding the usage of these newly discovered AMPs. The similarity between a newly discovered AMP and a chicken AMP in predicted 3D structures suggests a potential similarity in their modes of action [35], indicating a potential risk that a newly discovered AMP may lose its effect if some pathogens have already developed resistance to its reference chicken AMP [45]. However, the extent to which structural similarity implies similarity in modes of action remain untested. Properties other than the peptide structure itself, such as the distributions of the hydrophobic and positively charged residues, can all affect how an AMP works. Also, we note that none of our newly discovered AMPs have a 100% match in predicted 3D structures to the known chicken AMPs, and those predicted structures haven’t yet been confirmed through laboratory experimentation. Additionally, the structures of some AMPs may vary according to the surrounding microenvironment [46]. All of these cannot be observed through our current set of tests and warrant further study.

If cross-resistance to our newly discovered AMPs and known chicken AMPs does not exist in pathogens tested, then the novel AMPs with high similarities to known chicken AMPs in predicted structures but originating from other organisms may be prioritized as primary candidates for further translation into applications. These AMPs may have different evolutionary backgrounds from chicken AMPs, suggesting that many chicken pathogens may not have had enough opportunity to develop resistance against them. In the worst case, if cross-resistance to the newly discovered AMPs and known chicken AMPs does occur in some pathogens, then this work sets up a warning to the scientific community in the use of those AMPs in chicken farming.

Although the AMPs discovered in the present study were derived from putative precursor sequences from existing organisms, it still requires further investigation to determine whether those AMPs occur in nature. Due to the limitations of the *in silico* cleavage tool used, it is likely that some sequences, which may or may not be real precursors, were incorrectly cleaved, even when the resulting fragments displayed antimicrobial properties *in vitro*. It has been reported that the degradation products of some non-antimicrobial proteins with other biological functions exhibit antimicrobial activities [47], and some of the aforementioned protein fragments may belong to this category of AMPs. Further, we observed instances where the predicted AMPs originated from non-secreted proteins, such as GoGo1, which is a fragment of a transcriptional repressor. While these AMPs may be less likely to occur naturally, we still consider them as valuable discoveries. AMPs that do not exist in nature may possess unique advantages in that they may be “new” to most extant pathogens. Lastly, exploring the relationship between the antimicrobial properties of the novel AMPs and the originally annotated functions of their parent sequences in the database would be a promising direction for future research. This investigation may provide a better understanding of the full life cycle of these proteins or peptides.

As the UniProt database frequently updates with more protein sequence data from additional organisms, we expect the AMP mining workflow we present to continue being a valuable discovery and annotation engine. Future work can also be extended to additional peptide sequence databases (e.g., UniProtKB/TrEMBL, the unreviewed section of the UniProt database [28]). Additional work can be done by loosening the criteria in sequence filtering steps for more sequences. For example, sequences without any predicted signal peptides, which were all removed in Step 2 of our workflow (**Supplementary Figure S1**), can be secreted proteins or peptides reported in mature or incomplete precursor forms. Though it is out of the research scope of the current work, such sequences may hold value for further investigation. We expect bioinformatics workflows like the one reported here to continuously discover novel AMPs and to help the fight against MDR bacteria.

## Materials and methods

### Datasets

The present study involves four main datasets: 1) all eukaryotic entries from UniProtKB/Swiss-Prot [28], 2) entries annotated with AMP-related keywords in UniProtKB/Swiss-Prot [28], 3) known AMP sequences in AMP databases, and 4) known chicken AMP sequences.

All eukaryotic entries from the UniProtKB/Swiss-Prot database (2022_02 release) [28] were downloaded by using the query “(taxonomy_id:2759) AND (reviewed:true)”.

This set of sequences include 186,302 distinct sequences from 195,188 UniProt entries. The set of entries annotated with AMP-related keywords in the UniProtKB/Swiss-Prot database [28] includes 18,470 distinct sequences from 19,845 UniProt entries.

UniProt entries with their annotations containing any of the following 16 AMP-related keywords were downloaded: {antimicrobial, antibiotic, antibacterial, antiviral, antifungal, antimalarial, antiparasitic, anti-protist, anticancer, defense, defensin, cathelicidin, histatin, bacteriocin, microbicidal, fungicide} [18]. We note that 11,925 sequences from 12,229 UniProt entries in this set belong to the above eukaryotic sequence set.

The known AMP sequence set comprises 4,538 distinct AMP sequences. All AMP sequences were downloaded from two curated AMP databases: APD3 [9] on July 11, 2022, and Database of Anuran Defense Peptides (DADP, http://split4.pmfst.hr/dadp) [33] on December 6, 2018.

The known chicken AMP sequence set includes 22 sequences in total. These sequences were downloaded on October 14, 2022 from the APD3 database [9] by searching for source organism “*Gallus gallus*”.

### AMP mining workflow

The AMP mining workflow we report in the present study includes five steps, as shown in **Supplementary Figure S1**.

In Step 1, all eukaryotic protein sequences from the UniProtKB/Swiss-Prot database [28] were processed to generate candidate mature peptide sequences using the cleavage module of the AMP discovery pipeline rAMPage v1.0.1 [22]. The cleavage module of rAMPage is based on ProP v1.0c [32], a machine learning based tool for cleavage site prediction for eukaryotic protein sequences. The rAMPage cleavage module also provides post-processing of the ProP output [22]. ProP predicts all the cleavage sites but only annotates the corresponding signal peptides – it does not provide labels to mature peptide sequences and pro-sequences [22,32]. As a result, for each putative precursor sequence, the rAMPage cleavage module takes all cleaved pieces, excluding the predicted signal peptide, as well as all possible recombinations of non-adjacent cleaved pieces (with a maximum of three pieces within each recombination) as candidate mature peptide sequences [22]. Additionally, the rAMPage cleavage module removes candidate mature peptide sequences with lengths < 2 or > 200 aa, as AMPs are typically known to be shorter sequences [6]. The redundancy removal function of the rAMPage cleavage module was turned off in the present study, as the same putative AMP sequence from different UniProt entries were kept as different records (**Supplementary Figure S1**) for convenience of future investigations into their source organisms. However, we did consider them as the same putative AMP sequence when calculating the unique sequence counts presented.

In Step 2, sequences predicted to lack signal peptides were removed, as most AMPs are expected to be secreted [48].

In Step 3, the remaining candidate mature peptide sequences from Step 2 were passed through the imbalanced model of AMPlify v2.0.0 [19] for a preliminary filtering. Candidate mature peptide sequences predicted as AMPs by the AMPlify imbalanced model were passed on to the next step. Sequences with non-standard amino acids were also filtered out, as AMPlify does not assign those sequences with any predictions.

In Step 4, the predicted AMPs from Step 3 were then passed through the balanced model of AMPlify v2.0.0 [18] for a more precise filtering. Sequences that were predicted as AMPs by both AMPlify imbalanced and balanced models were collected as the final set of predicted AMPs.

In Step 5, all the predicted AMPs from Step 4 were checked against the known AMP sequence set as well as the annotations of the corresponding UniProt entries for novel putative AMPs. Predicted AMPs were only considered as novel putative AMPs if they were not in the known AMP sequence set and if their corresponding UniProt entry annotation did not have AMP-related keywords (see above under “Datasets”).

### Sequence and structural similarities between peptides

We used two measures of peptide similarity: sequence similarity and structural similarity.

The sequence similarity between two peptides was calculated as 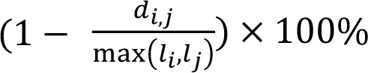, where *d_i,j_* is the edit distance and *l_i_*, *l_j_* are lengths of the peptide sequences. The structural similarity between two peptides was measured with TM-scores normalized by the average peptide sequence length [37] using TM-align [38]. TM-scores are between 0 and 1, with higher TM-scores indicating higher similarity between the two structures [37].

The sequence/structural similarity of a single peptide to a set of peptides was defined as the maximum of all sequence/structural similarity values calculated between that peptide and the peptides in the target set for comparison (i.e., the sequence/structural similarity of that peptide to its most similar target set peptide in their sequences/structures).

In this study, *in silico* predicted 3D structures were utilized for all peptide structural similarity comparisons. The 3D structures of peptides were predicted using ColabFold v1.4.0 [36]. ColabFold improves the prediction speed of AlphaFold2 [49] by combining it with MMseqs2 [50] homology search. For each peptide, five structures were generated by five AlphaFold2 models trained using different random seeds, with relaxations applied to the predicted structures utilizing gradient descent in the Amber force field [49,51]. The structures were then ranked by their average predicted local distance difference test (pLDDT) scores – a measure of the confidence of AlphaFold2 predictions. We chose the structure with the highest average pLDDT score for further analysis.

### Selecting putative AMPs for validation

Among all the novel putative AMPs discovered by the AMP mining workflow, we prioritized 397 short cationic peptides with AMPlify scores ≥ 10 (i.e., AMPlify probability scores ≥ 0.9) for selection, as shorter peptides are more cost-effective to synthesize [22]. We define short cationic peptides as those with lengths between 5 and 35 aa and net charge greater than 0. We note that AMPlify score is a prediction score reported by AMPlify, which is a log transformation of the AMPlify probability score *p_AMPlify_* as −10 log_10_ (1− *p_AMPlify_*), and here the AMPlify scores from the balanced model were taken for analysis.

First, the 3D structures of the 397 putative AMPs, together with the 22 known chicken AMPs, were predicted by ColabFold [36]. Next, the similarity (TM-score) of each putative AMP to the known chicken AMPs in predicted structures (i.e., similarity of the putative AMP to the most similar known chicken AMP in predicted 3D structures) was calculated using TM-align [38]. Finally, we ranked the peptides by their similarities to known chicken AMPs in predicted structures. The top 30 putative AMPs with highest TM-scores were prioritized for synthesis, as well as the bottom 10 with lowest TM-scores for comparison (**Supplementary Table S3**, **Supplementary Table S4**). Furthermore, the three reference chicken AMPs (i.e., the most similar known chicken AMP to a putative AMP in predicted 3D structures) for the top 30 putative AMPs: Chicken CATH-2 [39], Chicken CATH-3 [40], and Ovipin [41], were also sent for synthesis for comparison (**Supplementary Table S5**). We note that two of the top 30 putative AMPs could not be successfully synthesized, resulting in a final number of 38 putative AMPs in total proceeding to further *in vitro* validation.

### Antimicrobial susceptibility testing

To measure the antimicrobial activity of our selected putative AMPs *in vitro*, broth microdilution assays were conducted to determine the minimum inhibitory and minimum bactericidal concentrations (MICs and MBCs, respectively), as outlined by the Clinical and Laboratory Standards Institute (CLSI) [52]. The adaptations for testing cationic AMPs as described previously [53] were incorporated. The selected peptides were tested against laboratory isolates of *E. coli* 25922 and *S. aureus* 29213, which were purchased from the American Type Culture Collection (ATCC; Manassas, VA, USA). Bacteria from frozen stocks were streaked onto nonselective Columbia blood agar with 5% sheep blood (Oxoid) and incubated for 18–24 h at 37℃. To ensure the uniform colony health before the assay, 2–4 colonies were streaked onto a new agar plate and incubated for another 18–24 h at 37℃ on the next day. To create a standardized bacterial inoculum, isolated colonies were suspended in Mueller-Hinton Broth (MHB; Sigma-Aldrich, St. Louis. MO, USA). The suspension was adjusted to an optical density of 0.08–0.1 at 600 nm, which corresponds to a 0.5 McFarland standard of approximately 1–2 × 10^8^ CFU/mL (CFU: colony forming units). The inoculum was then diluted 1:250, resulting in a final concentration of 5±3 × 10^5^ CFU/mL. The target bacterial density was confirmed by measuring the total viability counts from the final tested inoculum.

The selected putative AMPs were purchased from and synthesized by GenScript (Piscataway, NJ, USA) in lyophilized format. The peptides were stored at -20℃, and were suspended in sterile ultrapure water prior to testing. A two-fold serial dilution of 1,280 to 2.5 μg/mL was prepared in sterile 96-well polypropylene microtitre plates (Greiner Bio-One #650261, Kremsmünster, Austria). Then, 100 μL of the standardized bacterial inoculum was added to each well, resulting in a final AMP testing range of 128 to 0.25 μg/mL. The MIC values were determined as the lowest peptide concentration in which no visible bacterial growth was observed after a 20–24 h incubation at 37℃. To determine the MBC values, well contents of the MIC and the two adjacent wells containing the two- and four-fold higher peptide concentrations were plated onto nonselective nutrient agar. The concentration in which 99.9% of the inoculum were killed after an incubation for 24 hours at 37℃ was reported as the MBC. In our tests, we used Ranatuerin-4 – a known AMP from the American bullfrog [54], and an in-house peptide [TKPKG]_3_ (OT15) as the positive and negative control peptides, respectively. We note that OT15 was truncated and derived from a negative control peptide [TKPKG]_4_ (OT20), not antimicrobial but sharing similar characteristics to AMPs, used in previous studies [55].

### Hemolysis assay

Hemolysis experiments were performed to evaluate the toxicity of the selected peptides to RBCs. Whole blood from healthy donor pigs was purchased from Lampire Biological Laboratories (Pipersville, PA, USA), and RBCs were washed and isolated by centrifugation using Roswell Park Memorial Institute (RPMI) medium (Life Technologies, Grand Island, NY, USA). Peptides were suspended and serially diluted from 1,280 to 10 μg/mL using RPMI medium in a 96-well polypropylene microtitre plate, and then combined with 100 μL of the 1% RBC solution, providing a final AMP testing range of 128 to 1 μg/mL. Plates were centrifuged after an incubation at 37℃ for 30–45 minutes, and 1/2 volume from each supernatant was transferred to a new 96-well plate. The absorbance of the wells was measured at 415 nm utilizing the Cytation 5 Cell Imaging Multimode Reader (BioTek, CA, USA), and the peptide concentration that lysed 50% of the RBCs (HC_50_) was used to determine the hemolytic activity. Absorbance readings from wells containing RBCs treated with 11 μL of a 2% Triton-X100 solution or RPMI medium (AMP solvent-only) were used to define 100% and 0% hemolysis, respectively. We note that all centrifugation steps were performed at 500× *g* for five minutes in an Allegra-6R centrifuge (Beckman Coulter, CA, USA).

## Author contributions

IB, RLW, LMNH, and CCH conceived the presented work with help from CL. MK conducted project administration. CL and IB designed the AMP mining workflow. CL collected and curated the datasets. CL implemented the AMP mining workflow with help from DS, DL, SIA, LC, and RLW. CL conducted peptide structure prediction with help from HE and AR. CL, DS, AY, RLW, and IB performed peptide selection. AS, DS, AR, and AY conducted antimicrobial susceptibility and hemolysis testing. CL, DS, AS, AR, AY, LC, RLW, and IB analyzed the results. CL drafted the manuscript, and all authors were involved in its revision. All authors read and approved the final manuscript.

