## Supplementary Material for "Mining the UniProtKB/Swiss-Prot database for antimicrobial peptides"

### Supplementary Figures

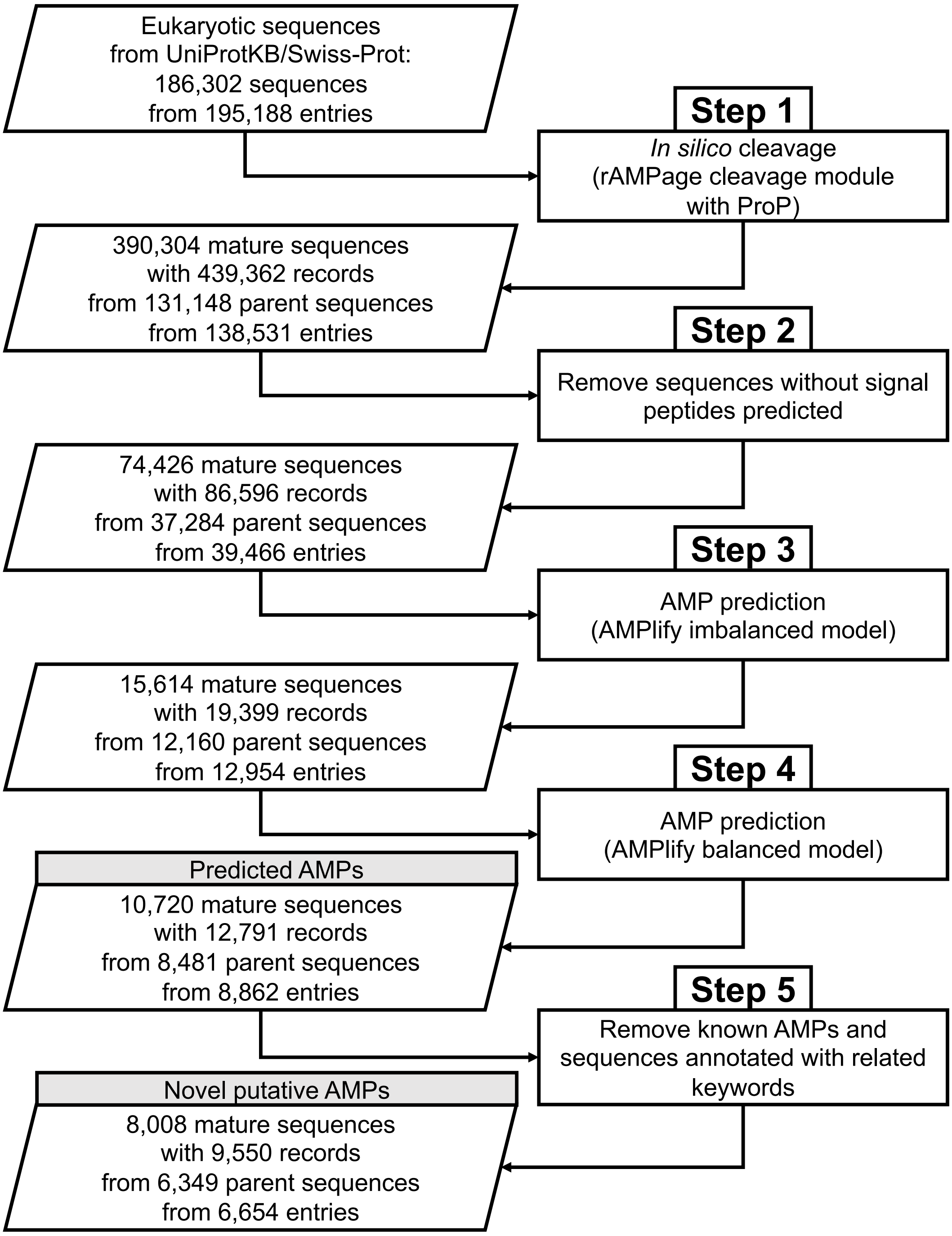


**Supplementary Figure S1: AMP mining workflow.** The AMP mining workflow utilizes the rAMPage [1] cleavage module with ProP [2] to cleave putative precursor sequences, and AMPlify [3,4] to predict AMP sequences. The numbers of candidate mature peptide sequences, candidate mature peptide sequence records, parent sequences, and UniProt entries, that remained at each step are reported. We note that two candidate mature peptide sequence records are considered distinct from each other if any of these three attributes are different: UniProt entry ID, position in the parent sequence, and candidate mature peptide sequence. All numbers presented in the workflow are non-redundant.

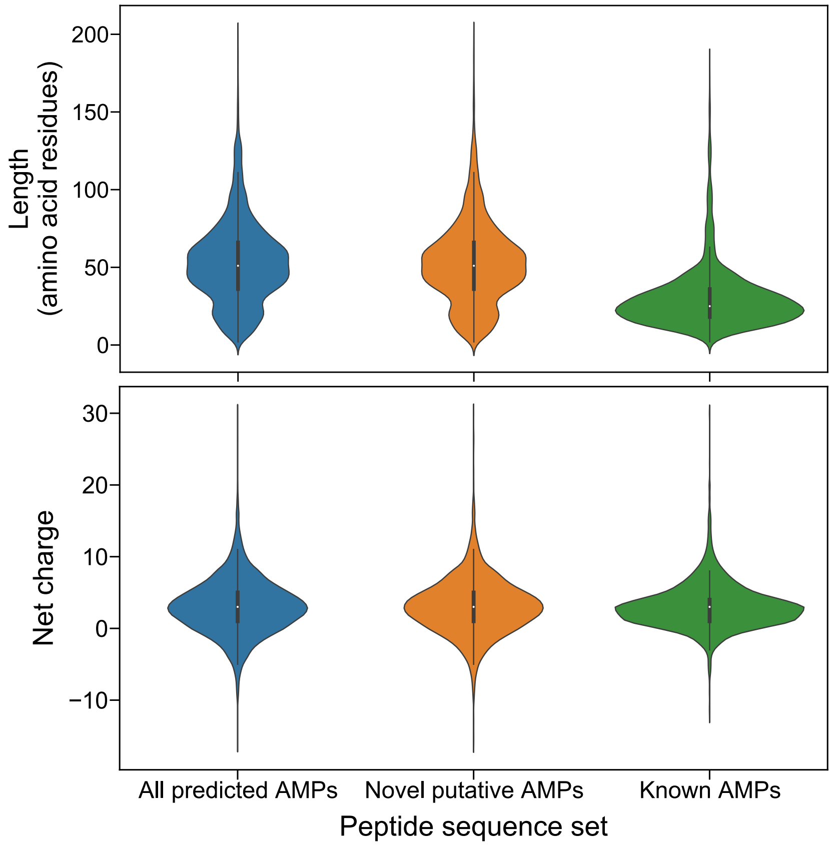


**Supplementary Figure S2: Length and net charge distributions of the AMPs predicted by AMPlify from the UniProtKB/Swiss-Prot database.** Length and net charge distributions were calculated for all 10,720 predicted AMPs as well as the 8,008 novel putative AMPs among them. The distributions of the 4,538 known AMP sequences from Antimicrobial Peptide Database (APD3) [5] and Database of Anuran Defense Peptides (DADP) [6] were plotted alongside for comparison. Mean (μ) and standard deviation (σ) values of each distribution are as follows: all predicted AMPs (length: μ = 52.83 aa, σ = 26.59 aa; net charge: μ= 3.02, σ = 3.90), novel putative AMPs (length: μ = 52.49 aa, σ = 26.49 aa; net charge: μ = 3.04, σ = 3.88), and known AMPs (length: μ = 30.21 aa, σ = 20.28 aa; net charge: μ = 3.05, σ = 3.10).

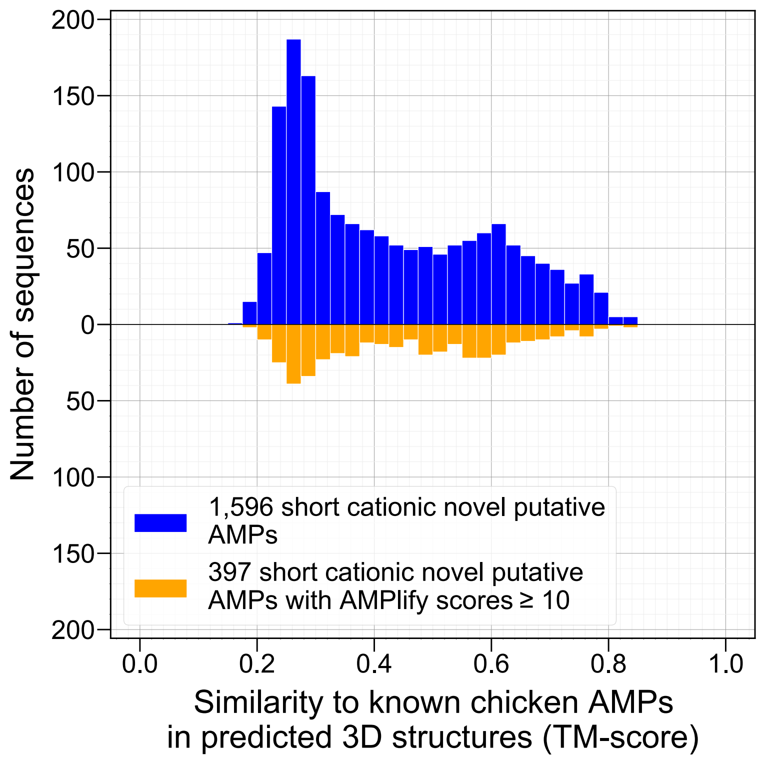


**Supplementary Figure S3: Similarity distributions of the short cationic novel putative AMPs mined from the UniProtKB/Swiss-Prot database to known chicken AMPs in predicted three-dimensional structures.** The similarity distribution of 1,596 short cationic novel putative AMPs to known chicken AMPs in predicted three-dimensional (3D) structures holds a mean of 0.4266 and a standard deviation of 0.1690. Among all short cationic novel putative AMPs, 397 of them are with AMPlify scores ≥ 10 (i.e., AMPlify probability scores ≥ 0.9). The similarity distribution of these 397 putative AMPs to known chicken AMPs in predicted 3D structures holds a mean of 0.4434 and a standard deviation of 0.1611. The similarity of each putative AMP to known chicken AMPs in predicted 3D structures (TM-score) was considered as the similarity of the putative AMP to the most similar known chicken AMP in predicted 3D structures (i.e., reference chicken AMP) from Antimicrobial Peptide Database (APD3) [5], based on which the distributions were plotted.

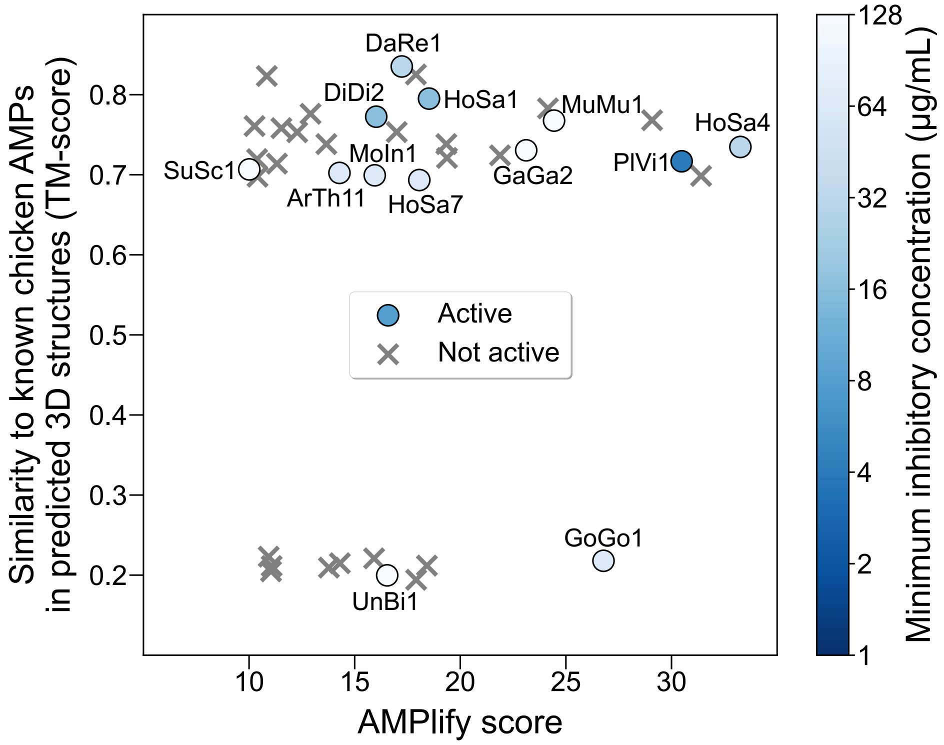


**Supplementary Figure S4: Visualization of antimicrobial activity of the 38 tested putative AMPs with respect to AMPlify scores and associated similarities to known chicken AMPs in predicted three-dimensional structures.** Peptides without any observable antimicrobial activity are presented as grey crosses, and the active peptides are presented in blue dots with their names annotated. Dots with darker colors indicate stronger antimicrobial activity against *Escherichia coli* ATCC 25922, determined by the lowest minimum inhibitory concentration (MIC) value of each peptide against the strain. AMPlify scores from the balanced model were used for visualization. The similarity of each tested peptide to known chicken AMPs in predicted three-dimensional (3D) structures (TM-score) was considered as the similarity of that peptide to the most similar known chicken AMP in predicted 3D structures (i.e., reference chicken AMP) from Antimicrobial Peptide Database (APD3) [5].

### Supplementary Tables

**Supplementary Table S1: Performance comparison among different tools on the balanced test set.** Values of accuracy (acc), sensitivity (sens), specificity (spec), F1 score (F1) and area under the receiver operating characteristic curve (AUROC) are presented in percentages.

| Tool | Model | Acc | Sens | Spec | F1 | AUROC |
|---|---|---|---|---|---|---|
| iAMPpred [7] | original[a] | 74.01 | 87.90 | 60.12 | 77.18 | 80.70 |
| iAMP-2L [8] | original[a] | 77.96 | 88.26 | 67.66 | 80.02 | —* |
| AMP Scanner Vr.2 [9] | original[a] | 78.50 | 90.66 | 66.35 | 80.83 | 88.33 |
| AMPlify [3,4] | balanced | **93.71** | 92.93 | 94.49 | **93.66** | **98.37** |
| | imbalanced | 89.22 | **94.37** | 84.07 | 89.75 | 96.88 |
| | balanced + imbalanced[b] | 93.59 | 91.50 | **95.69** | 93.46 | —* |

[a] Models presented in the referenced papers, which are available through online servers.

[b] Only sequences predicted as AMPs by both models are determined to be AMPs.

* AUROC values were not calculated for models which only output predicted labels instead of probabilities.

**Supplementary Table S2: Performance comparison among different tools on the imbalanced test set.** Values of accuracy (acc), sensitivity (sens), specificity (spec), F1 score (F1) and area under the receiver operating characteristic curve (AUROC) are presented in percentages. We note that the online server of iAMP-2L [8] was down at the time this analysis was done (November 11, 2022), so its results were not compared here.

| Tool | Model | Acc | Sens | Spec | F1 | AUROC |
|---|---|---|---|---|---|---|
| iAMPpred [7] | original[a] | 14.29 | 87.90 | 11.90 | 6.07 | 55.54 |
| AMP Scanner Vr.2 [9] | original[a] | 71.26 | 90.66 | 70.63 | 16.57 | 91.44 |
| AMPlify [3,4] | balanced | 95.97 | 92.93 | 96.06 | 59.19 | 98.06 |
| | imbalanced | 98.94 | **94.37** | 99.09 | 84.87 | **99.54** |
| | balanced + imbalanced[b] | **99.31** | 91.50 | **99.56** | **89.25** | —[*] |

[a] Models presented in the referenced papers, which are available through online servers.

[b] Only sequences predicted as AMPs by both models are determined to be AMPs.

[*] AUROC values were not calculated for models which only output predicted labels instead of probabilities.

**Supplementary Table S3: Characteristics of the 40 putative AMP sequences mined from the UniProtKB/Swiss-Prot database that have been prioritized for synthesis.** A total number of 397 short cationic novel putative AMPs with AMPlify scores ≥ 10 (i.e., AMPlify probability scores ≥ 0.9) were compared with known chicken AMPs for their similarities in predicted three-dimensional (3D) structures. The top section shows the 30 most similar putative AMPs to the known chicken AMPs in predicted 3D structures as measured by TM-scores, while the bottom section shows the 10 least similar ones to the known chicken AMPs in predicted 3D structures.

| **Peptide name** | **Sequence** | **# aa** | **Net charge[a]** | **Molecular weight (Da)** | **AMPlify score[b]** | **Sequence similarity to known AMPs[c] (%)** | **Similarity to known chicken AMPs in predicted 3D structures[d] (TM-score)** | **Reference chicken AMP[e]** |
|---|---|---|---|---|---|---|---|---|
| DaRe1 | Sequence information is embargoed. | 25 | 5 | 3027.63 | 17.23 | 40.00 | 0.8352 | Chicken CATH-2 |
| ScPo1 | | 32 | 2 | 4138.74 | 17.90 | 37.50 | 0.8252 | Chicken CATH-2 |
| ArTh10 | | 31 | 4 | 3716.39 | 10.84 | 35.48 | 0.8233 | Chicken CATH-2 |
| HoSa1 | | 26 | 6 | 2921.51 | 18.52 | 38.46 | 0.7951 | Chicken CATH-2 |
| HoSa2 | | 22 | 6 | 2664.21 | 24.15 | 33.33 | 0.7829 | Chicken CATH-3 |
| DiVi1 | | 29 | 5 | 3315.83 | 12.91 | 36.36 | 0.7761 | Chicken CATH-3 |
| DiDi2 | | 30 | 6 | 3662.31 | 16.02 | 33.33 | 0.7725 | Chicken CATH-2 |
| DiDi3[*] | | 25 | 1 | 2650.45 | 30.78 | 44.00 | 0.7724 | Chicken |

| | | | | | | | | |
|---|---|---|---|---|---|---|---|---|
| | | | | | | | | CATH-3 |
| LySt2 | | 31 | 5 | 3652.26 | 29.08 | 35.48 | 0.7681 | Chicken CATH-3 |
| MuMu1 | | 35 | 8 | 4135.81 | 24.44 | 34.29 | 0.7676 | Chicken CATH-2 |
| CaGl1 | | 30 | 1 | 3759.20 | 10.26 | 33.33 | 0.7608 | Chicken CATH-3 |
| HoSa3 | | 22 | 3 | 2653.16 | 11.53 | 40.91 | 0.7582 | Chicken CATH-3 |
| BoMo2 | | 24 | 3 | 2790.11 | 16.99 | 33.33 | 0.7533 | Chicken CATH-2 |
| ScPo2 | | 22 | 4 | 2722.25 | 12.28 | 36.36 | 0.7531 | Chicken CATH-3 |
| DiDi1 | | 23 | 2 | 2819.51 | 19.34 | 34.78 | 0.7383 | Chicken CATH-3 |
| RaNo1 | | 34 | 5 | 4203.07 | 13.66 | 35.29 | 0.7383 | Chicken CATH-3 |
| HoSa4 | | 20 | 5 | 2682.28 | 33.26 | 35.00 | 0.7347 | Chicken CATH-3 |
| GaGa2 | | 11 | 5 | 1380.75 | 23.12 | 45.45 | 0.7305 | Ovipin |
| MuMu2* | | 22 | 2 | 2345.04 | 10.44 | 40.91 | 0.7250 | Chicken CATH-3 |
| RaNo2 | | 24 | 1 | 2871.37 | 21.88 | 29.17 | 0.7242 | Chicken CATH-2 |
| HoSa5 | | 25 | 7 | 3111.67 | 19.37 | 32.00 | 0.7210 | Chicken |

| | | | | | | | | |
|---|---|---|---|---|---|---|---|---|
| | | | | | | | | CATH-3 |
| EiHe1 | | 31 | 2 | 3704.47 | 10.38 | 38.71 | 0.7195 | Chicken CATH-3 |
| PlVi1 | | 19 | 5 | 2390.01 | 30.48 | 40.00 | 0.7168 | Chicken CATH-3 |
| MuMu3 | | 34 | 3 | 4130.92 | 11.32 | 32.35 | 0.7139 | Chicken CATH-2 |
| SuSc1 | | 23 | 6 | 2787.27 | 10.01 | 34.78 | 0.7067 | Chicken CATH-3 |
| ArTh11 | | 22 | 9 | 2769.34 | 14.28 | 36.36 | 0.7023 | Chicken CATH-3 |
| MoIn1 | | 19 | 2 | 2281.77 | 15.95 | 31.58 | 0.6993 | Chicken CATH-2 |
| HoSa6 | | 11 | 5 | 1333.69 | 31.40 | 42.11 | 0.6985 | Ovipin |
| CaGl2 | | 20 | 3 | 2485.11 | 10.40 | 50.00 | 0.6976 | Chicken CATH-3 |
| HoSa7 | | 35 | 3 | 3926.73 | 18.06 | 28.57 | 0.6934 | Chicken CATH-3 |
| GaGa4 | | 20 | 3 | 2181.49 | 10.93 | 35.00 | 0.2230 | OvoDBβ |
| OrSa6 | | 12 | 4 | 1321.52 | 15.92 | 37.50 | 0.2209 | OvoDBβ |
| GoGo1 | | 22 | 11 | 2457.84 | 26.78 | 44.12 | 0.2179 | CHB1 |
| HoSa8 | | 6 | 3 | 872.04 | 14.30 | 42.86 | 0.2148 | Ovipin |
| HoSa9 | | 23 | 3 | 2624.99 | 18.42 | 34.78 | 0.2119 | CHB2 |
| DrMe6 | | 9 | 3 | 1037.23 | 11.07 | 37.50 | 0.2113 | Chicken Heterophil |

| | | | | | | | | |
|---|---|---|---|---|---|---|---|---|
| | [redacted] | | | | | | | Peptide 2 |
| MuMu4 | [redacted] | 17 | 2 | 1734.07 | 13.78 | 47.06 | 0.2094 | OvoDBβ |
| HoSa10 | [redacted] | 7 | 2 | 775.95 | 11.04 | 41.67 | 0.2047 | OvoDBβ |
| UnBi1 | [redacted] | 8 | 2 | 957.19 | 16.54 | 41.67 | 0.1997 | Ovipin |
| ScPo3 | [redacted] | 6 | 2 | 736.87 | 17.91 | 42.86 | 0.1942 | Ovipin |

[a] Net charge at pH = 7.

[b] AMPlify scores from the balanced model. AMPlify scores range from 0 to 80; sequences with AMPlify scores > 3.01 (i.e., AMPlify probability scores > 0.5) are predicted as AMPs.

[c] Sequence similarity to the most similar known AMP sequence from Antimicrobial Peptide Database (APD3, downloaded on July 11, 2022) [5] and Database of Anuran Defense Peptides (DADP, downloaded on December 6, 2018) [6].

[d] Similarity to the most similar known chicken AMP in predicted 3D structures (i.e., reference chicken AMP) from APD3 [5] (downloaded on October 14, 2022). 3D structures of the peptides were predicted by ColabFold [10,11], and the similarity between two peptides in predicted 3D structures were evaluated by TM-score [12] normalized by the average peptide sequence length using TM-align [13].

[e] The most similar known chicken AMP in predicted 3D structures.

[*] DiDi3 and MuMu2 were not successfully synthesized and were excluded for further tests.

**Supplementary Table S4: Overview of the 40 putative AMP sequences mined from the UniProtKB/Swiss-Prot database that have been prioritized for synthesis regarding their corresponding parent sequence information in the database.** This table supplements Supplementary Table S3 with information about the 40 putative AMP sequences regarding their corresponding parent sequences. The top section shows the 30 most similar putative AMPs to the known chicken AMPs in predicted three-dimensional (3D) structures as measured by TM-scores, while the bottom section shows the 10 least similar ones to the known chicken AMPs in predicted 3D structures.

| Putative AMP name | Position in parent sequence[a] | UniProt entry ID | Source organism | Source organism category[b] |
|---|---|---|---|---|
| DaRe1 | 17 – 41 | B0S8I0 | *Danio rerio* | Others |
| ScPo1 | 20 – 51 | Q9Y7K8 | *Schizosaccharomyces pombe* | Others |
| ArTh10 | [29 – 45] + [356 – 369] | Q7Y223 | *Arabidopsis thaliana* | Plant |
| HoSa1 | 299 – 324 | Q8NH93 | *Homo sapiens* | Mammal |
| HoSa2 | 34 – 55 | Q8IYJ2 | *Homo sapiens* | Mammal |
| DiVi1 | 983 – 1011 | Q24702 | *Dictyocaulus viviparus* | Others |
| DiDi2 | 38 – 67 | Q54GV3 | *Dictyostelium discoideum* | Others |
| DiDi3* | 145 – 169 | Q54UP0 | *Dictyostelium discoideum* | Others |
| LySt2 | [187 – 199] + [223 – 225] + [292 – 306] | P19802 | *Lymnaea stagnalis* | Others |
| MuMu1 | 16 – 50 | P09925 | *Mus musculus* | Mammal |
| CaGl1 | 31 – 60 | Q6FWE8 | *Candida glabrata* | Others |
| HoSa3 | 145 – 166 | Q9H3J6 | *Homo sapiens* | Mammal |
| BoMo2 | 44 – 67 | P82003 | *Bombyx mori* | Insect |
| ScPo2 | 36 – 57 | O74430 | *Schizosaccharomyces pombe* | Others |
| DiDi1 | 539 – 561 | Q54TM2 | *Dictyostelium discoideum* | Others |
| RaNo1 | 194 – 227 | P09320 | *Rattus norvegicus* | Mammal |

| | | | | |
|---|---|---|---|---|
| HoSa4 | 39 – 58 | A8MTL0 | *Homo sapiens* | Mammal |
| GaGa2 | 23 – 33 | P10039 | *Gallus gallus* | Others |
| MuMu2* | 456 – 477 | Q2TB54 | *Mus musculus* | Mammal |
| RaNo2 | 110 – 133 | Q5U2T1 | *Rattus norvegicus* | Mammal |
| HoSa5 | [23 – 38] + [189 – 197] | O75596 | *Homo sapiens* | Mammal |
| EiHe1 | 32 – 62 | A0A291NUI5 | *Eidolon helvum* | Mammal |
| PlVi1 | 99 – 117 | P0CV63 | *Plasmopara viticola* | Others |
| MuMu3 | 194 – 227 | O35256 | *Mus musculus* | Mammal |
| SuSc1 | 22 – 44 | P53366,<br>O62827 | *Sus scrofa*,<br>*Bos taurus* | Mammal |
| ArTh11 | 28 – 49 | Q6NKN8 | *Arabidopsis thaliana* | Plant |
| MoIn1 | 1861 – 1879 | Q09WW0 | *Morus indica* | Plant |
| HoSa6 | 23 – 33 | P24821,<br>Q29116 | *Homo sapiens*,<br>*Sus scrofa* | Mammal |
| CaGl2 | 29 – 48 | P05040 | *Candida glabrata* | Others |
| HoSa7 | 75 – 109 | Q99678 | *Homo sapiens* | Mammal |
| GaGa4 | [25 – 29] + [286 – 300] | Q92080 | *Gallus gallus* | Others |
| OrSa6 | 25 – 36 | Q5VRI5 | *Oryza sativa subsp. japonica* | Plant |
| GoGo1 | 744 – 765 | A1YF22,<br>A1YG99,<br>A2T7S4,<br>A2T771,<br>Q9UKY1 | *Gorilla gorilla gorilla*,<br>*Pan paniscus*,<br>*Pongo pygmaeus*,<br>*Pan troglodytes*,<br>*Homo sapiens* | Mammal |
| HoSa8 | 200 – 205 | Q9BXP8 | *Homo sapiens* | Mammal |
| HoSa9 | 562 – 584 | Q68DV7 | *Homo sapiens* | Mammal |

| | | | | |
|---|---|---|---|---|
| DrMe6 | 17 – 25 | O46201 | *Drosophila melanogaster* | Insect |
| MuMu4 | 469 – 485 | Q9Z0L3 | *Mus musculus* | Mammal |
| HoSa10 | 286 – 292 | Q86YB7 | *Homo sapiens* | Mammal |
| UnBi1 | 10 – 17 | C0HKK6 | *Unedogemmula bisaya* | Others |
| ScPo3 | 24 – 29 | O42663 | *Schizosaccharomyces pombe* | Others |

[a] Position of the putative mature AMP sequence in the corresponding parent sequence. For a putative AMP sequence that is the recombination of multiple cleaved peptide sequences, the positions of those cleaved peptide sequences are presented in brackets with plus signs connecting them to each other.

[b] Source organisms are classified into five categories: amphibian, plant, insect, mammal, and others.

[*] DiDi3 and MuMu2 were not successfully synthesized and were excluded for further tests.

**Supplementary Table S5: Seven reference chicken AMP sequences for the 40 putative AMPs mined from the UniProtKB/Swiss-Prot database and prioritized for synthesis.** The AMP sequences listed in this table are the seven reference chicken AMPs for the 40 putative AMPs listed in Supplementary Table S3. Only the three reference chicken AMPs for the top 30 putative AMPs which share highest similarities to known chicken AMPs in predicted three-dimensional (3D) structures were prioritized for synthesis and further tests (i.e., Chicken CATH-2, Chicken CATH-3, and Ovipin).

| APD3 ID | Peptide name | Sequence | # aa | Net charge[a] | Molecular weight (Da) |
|---|---|---|---|---|---|
| AP00548 | Chicken CATH-2 [14] | RFGRFLRKIRRFRPKVTITI QGSARFG | 27 | 9 | 3264.92 |
| AP00613 | Chicken CATH-3 [15] | RVKRFWPLVPVAINTVAA GINLYKAIRRK | 29 | 7 | 3351.09 |
| AP03457 | Ovipin [16] | YVSPVAIVKGLNIPL | 15 | 1 | 1582.95 |
| AP03014 | OvoDBβ [17] | QSKKCCGRCSSRMCTKRE KEEHTEDCRGSFCCLTHR KKK | 39 | 7 | 4609.37 |
| AP02878 | CHB1 [18] | VLSAADKNNVKGIFTKIA GHAEEYGAETLERMFTTY PPTKTY | 42 | 0 | 4664.27 |
| AP02879 | CHB2 [18] | LTAEDKKLIQQAWEKAAS HQEEFGAEALTRMFTTYP QTKTY | 41 | -1 | 4762.33 |
| AP00265 | Chicken Heterophil Peptide 2 [19] | GRKSDCFRKNGFCAFLKC PYLTLISGLCSFHLC | 33 | 4 | 3729.47 |

[a] Net charge at pH = 7.

**Supplementary Table S6: Antimicrobial susceptibility testing and hemolysis experiment results of the 38 successfully synthesized putative AMPs mined from the UniProtKB/Swiss-Prot database and tested *in vitro*.** Peptides were tested for their antimicrobial activity against *Escherichia coli* ATCC 25922 and *Staphylococcus aureus* ATCC 29213 for their minimum inhibitory concentration (MIC) and minimum bactericidal concentration (MBC) values. Porcine red blood cells (RBCs) were used to test the hemolytic activity of the selected peptides for their hemolytic concentration ($HC_{50}$) values. Data is presented as the lowest effective peptide concentration range (µg/mL) observed in three independent experiments performed in duplicate, with one maximum data point and one minimum data point dropped for each measurement. The top section shows results of the 28 successfully synthesized peptides among the top 30 peptides in similarities to known chicken AMPs in predicted three-dimensional (3D) structures as measured by TM-scores, while the second section shows results of the 10 least similar ones to known chicken AMPs in predicted 3D structures. Results of the three reference chicken AMPs (Chicken CATH-2, Chicken CATH-3, and Ovipin) for the top 30 peptides are listed in the third section for comparison. The control peptides in the bottom section includes: a positive control peptide Ranatuerin-4 [20] and a negative control peptide OT15.

| Peptide name | Antimicrobial susceptibility testing | | | | Hemolysis testing[a] |
|---|---|---|---|---|---|
| | *E. coli* ATCC 25922 | | *S. aureus* ATCC 29213 | | Porcine RBCs |
| | MIC (µg/mL) | MBC (µg/mL) | MIC (µg/mL) | MBC (µg/mL) | $HC_{50}$ (µg/mL) |
| DaRe1 | 32 – 64 | 32 – 64 | >128 | >128 | >128 |
| ScPo1 | >128 | >128 | >128 | >128 | — |
| ArTh10 | >128 | >128 | >128 | >128 | — |
| HoSa1 | 16 | 16 – 32 | 128 | —* | >128 |
| HoSa2 | >128 | >128 | >128 | >128 | — |
| DiVi1 | >128 | >128 | >128 | >128 | — |
| DiDi2 | 16 | 16 | >128 | >128 | >128 |
| LySt2 | >128 | >128 | >128 | >128 | — |
| MuMu1 | ≥128 | —* | >128 | >128 | — |
| CaGl1 | >128 | >128 | >128 | >128 | — |

| | | | | | |
|---|---|---|---|---|---|
| HoSa3 | >128 | >128 | >128 | >128 | — |
| BoMo2 | >128 | >128 | >128 | >128 | — |
| ScPo2 | >128 | >128 | >128 | >128 | — |
| DiDi1 | >128 | >128 | >128 | >128 | — |
| RaNo1 | >128 | >128 | >128 | >128 | — |
| HoSa4 | 32 | 32 | 64 – 128 | —* | >128 |
| GaGa2 | 128 | —* | >128 | >128 | >128 |
| RaNo2 | >128 | >128 | >128 | >128 | — |
| HoSa5 | >128 | >128 | >128 | >128 | — |
| EiHe1 | >128 | >128 | >128 | >128 | — |
| PlVi1 | 4 – 8 | 8 | 32 | 32 | >128 |
| MuMu3 | >128 | >128 | >128 | >128 | — |
| SuSc1 | 128 | —* | >128 | >128 | >128 |
| ArTh11 | 64 – 128 | —* | >128 | >128 | >128 |
| MoIn1 | 64 – >128 | —* | >128 | >128 | — |
| HoSa6 | >128 | >128 | >128 | >128 | — |
| CaGl2 | >128 | >128 | >128 | >128 | — |
| HoSa7 | 64 – >128 | —* | >128 | >128 | — |
| GaGa4 | >128 | >128 | >128 | >128 | — |
| OrSa6 | >128 | >128 | >128 | >128 | — |
| GoGo1 | 64 | —* | >128 | >128 | >128 |
| HoSa8 | >128 | >128 | >128 | >128 | — |
| HoSa9 | >128 | >128 | >128 | >128 | — |
| DrMe6 | >128 | >128 | >128 | >128 | — |
| MuMu4 | >128 | >128 | >128 | >128 | — |

| | | | | | |
|---|---|---|---|---|---|
| HoSa10 | >128 | >128 | >128 | >128 | — |
| UnBi1 | 128 | —* | >128 | >128 | >128 |
| ScPo3 | >128 | >128 | >128 | >128 | >128 |
| Chicken CATH-2 | 8 – 16 | 8 – 16 | 32 | 32 | >128 |
| Chicken CATH-3 | 2 – 4 | 2 – 4 | 2 | 2 – 4 | >128 |
| Ovipin | >128 | >128 | >128 | >128 | >128 |
| Ranatuerin-4 | 4 | 4 | 1 – 2 | 2 | 16 |
| OT15[b] | >128 | >128 | >128 | >128 | >128 |

[a] Hemolysis experiments were not performed for putative AMPs that did not show any antimicrobial activity (MIC > 128 µg/mL) in at least two repeats for each bacterial strain tested.

[b] OT15 (TKPKGTKPKGTKPKG) is a truncated form of a negative control peptide OT20 [21] used in previous studies.

* MBC values were not tested for experiments revealing antimicrobial activity but with high MIC values of ≥ 64 µg/mL.

'—' = not tested.
